# Automated classification of signal sources in mesoscale calcium imaging

**DOI:** 10.1101/2021.02.23.432573

**Authors:** Brian R. Mullen, Sydney C. Weiser, Desiderio Ascencio, James B. Ackman

## Abstract

Functional imaging of neural cell populations is critical for mapping intra− and inter−regional network dynamics across the neocortex. Recently we showed that an unsupervised machine learning decomposition of densely sampled recordings of cortical calcium dynamics results in a collection of components comprised of neuronal signal sources distinct from optical, movement, and vascular artifacts. Here we build a supervised learning classifier that automatically separates neural activity and artifact components, using a set of extracted spatial and temporal metrics that characterize the respective components. We demonstrate that the performance of the machine classifier matches human identification of signal components in novel data sets. Further, we analyze control data recorded in glial cell reporter and non−fluorescent mouse lines that validates human and machine identification of functional component class. This combined workflow of data−driven video decomposition and machine classification of signal sources will aid robust and scalable mapping of complex cerebral dynamics.

## Introduction

A combination of advances in genetic engineering, parallel signal acquisition, and computation have enabled tremendously increased amounts of time series data concerning the function of neural systems ^1^. As a result, development of analytic tools to aid in the extraction of meaning from these large datasets is being undertaken. Implementation of unsupervised and supervised machine learning strategies can aid in extraction and interpretation of signals from complex time series data increasingly common in systems neuroscience. Machine learning techniques have been utilized to filter experimental artifacts, perform signal de−mixing, and understand brain−wide functional associations in data recorded with techniques such as fMRI and EEG ^2−4^. While analytic pipelines exist for these datasets, other types of functional recordings have their own advantages, as well as experimental challenges.

Mesoscale functional imaging across the dorsal cerebral cortical surface is used to map and understand dynamics that bridge inter−regional (macroscale) and inter−cellular (microscale) datasets. Large calcium imaging datasets have been popularized due to advances in mouse transgenics and the ability to record calcium flux activity simultaneously from many neurons ^5^. Analyzing population dynamics within and between regional networks in high supracellular resolution recordings can further our understanding of information processing in the brain. Mesoscale calcium imaging has been used to investigate patterns of neural activity that assist in self−organization of the developing brain ^6, 7^. Functional imaging in behaving mice has been utilized to identify new cortical subregions ^8^. Additionally, wide−field calcium imaging of the neocortex during electrode stimulation has helped map information flow between distant cerebral subregions ^9^.

Determination of neural signals from calcium imaging sessions in unanesthetized, behaving mice is an essential, but challenging task due to numerous confounding signal sources. Vascular artifacts are commonly seen due to vasodynamics and the resulting changes in blood flow to meet the energy demands of surrounding tissue. Fluid exchange between vascular and neural tissue causes cortical hemodynamics, which results in region specific changes of optical properties among cerebral lobes ^10^. Further, during the experimental preparation the skull is typically fixed to a specific location, however slight brain movements occur within the cranium that influence the recordings. Any optical property differences that originate from the experimental preparation may be highlighted in the dataset as signal due to changes in tissue contrast.

Principled exploration and identification of each signal source is necessary to reduce the contamination resulting from these physiological dynamics. Independent component analysis (ICA) is a nonparametric unsupervised machine learning technique that we utilize to identify each signal source in densely sampled (5.5 million pixels per frame) calcium imaging videos based on their spatially co−activating pixels and temporal properties ^11^. The global mean timecourse was initially subtracted and stored, thereby zeroing out each frame and allowing ICA to decompose each signal distinct from global effects. The decomposition results in hundreds of neural source components per hemisphere that are distinctly de−mixed from from artifact source signals.

In this study, we determine necessary conditions for identifying neural signals in mesoscale cortical calcium flux videos and create a machine learning classifier to isolate neural activity for analysis. We show that a combination of morphological and temporal metrics can characterize the signal components that maximize statistical independence in the video and we build a classifier that automatically selects for neural components. Our concurrent analysis of control wide−field imaging data corroborates the identification of artifact signal sources and gives insight into the structure of neuronal calcium dynamics across neocortex.

## Results

ICA was successfully used to extract components from mesoscale imaging data across the dorsal cortex of postnatal day 21 unanesthetized, pan−neuronal GCaMP6s expressing mice. Each component has a spatially decomposed eigenvector with its corresponding relative intensity temporal fluctuations. Similar data was recorded and processed in three sets of age matched control mice: cx3cr1 GFP (microglia; mGFP), adhl1 GFP (astrocyte; aGFP), and the non−transgenic C57/black 6 (Bl6) mice.

### Spatiotemporal metrics from components

Inspection of each set of experimental components resulted in the human classification as neural or artifact(fig. 1a). The artifacts were further distinguished between vascular and other for descriptive purposes. Neural components typically have globular spatial representation with highly dynamic properties. Vascular artifact components can be easily identified by the vascular−like spatial representation, which are also temporally dynamic. Other artifact components that are commonly seen in the components are movement or preparation artifacts. These typically have a diffuse spatial representation with smaller or sparse temporal activations. We manually scored each component in the dataset as an artifact (vascular or other) or neural component [(fig. 1b). From all the GCaMP experiments, on average 73.5 *±* 5.9% of the components were identified as neuronal, where the remaining 26.5 *±* 6.3% were artifact (vascular: 8.7 *±* 2.7%; other: 17.6 *±* 7.1%). GCaMP mice had substantially higher numbers of neural components compared to the controls, resulting in four times as many as the GFP mouse lines and six times the number in Bl6 mice (mean number of neural components GCaMP: 235, mGFP:62, aGFP:54, Bl6: 39).

**Figure 1:**
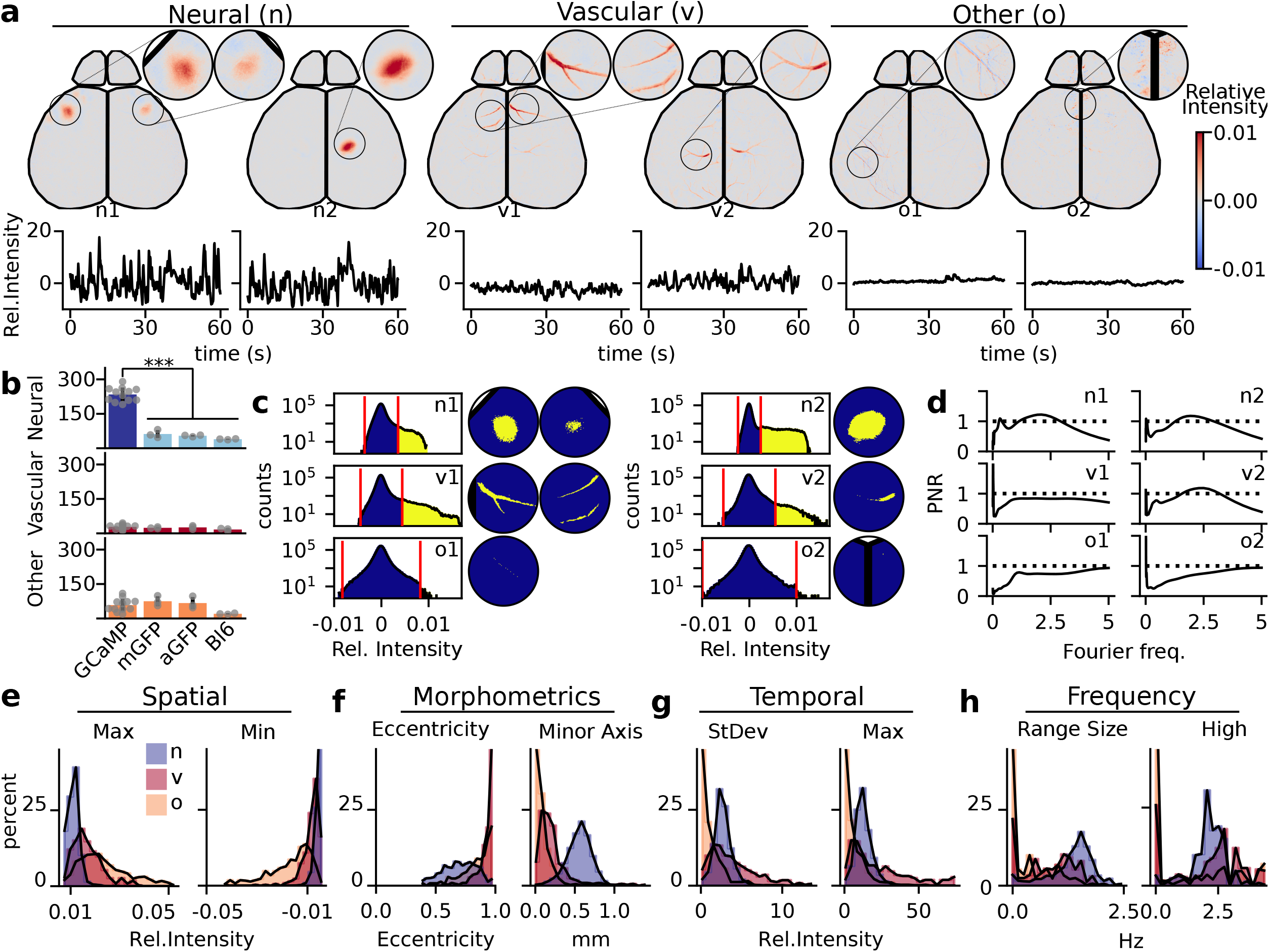
Class identity cannot be established by any individual extracted feature. (a) Examples of independent components of neural (n) signal, vascular (v) artifacts, and other (o) artifacts. Components are defined by both the spatial representation (eigenvector) and its temporal fluctuations. Circular windows magnify key portions of the eigenvector. Eigenvector values represented by colormap from blue to red. Temporal representation is in relative intensity (black time course under the eigenvector), only 1 minute of the full 20 minutes are shown. (b) A comparison of the number of neural signal (GCaMP: dark blue; controls: light blue) and the artifact components (vascular: red; other : orange) with each animal shown (GCaMP components: N=12 animals, n=3851; mGFP components: N=3, n = 484; aGFP components: N=3, n = 442; WT components: N=3, n=229). (c) Examples of binarization of the eigenvector. Histogram shows the full distribution of eigenvector values. The dynamic threshold method to generate binarized masks was used to identify the high eigenvector signal pixels (yellow) against the gaussian background (blue). Windowed spatial representation shows binarization on the key portions of the eigenvector. (d) Examples of neural and artifact wavelet analysis shown in the to signal power−noise ratio (PNR) plots. 95% red−noise cutoff was used to create signal to noise ratio (black dashed lines). (e) Histograms of example spatial metrics derived from GCaMP eigenvector values, (f) morphometrics from the shape of the binarized primary region, (g) temporal metrics derived from relative temporal intensities, (h) frequency metrics derived from the PNR.

We extracted spatial and morphological metrics of the neural and artifact components to characterize spatial feature differences (fig. 1c). From each component’s spatial eigenvector we can pull general spatial intensity metrics like global minimum and maximum. Further, the largest eigenvector values correspond to the regions that have the most dynamic change after the video rebuild. Given that the shapes of the high pixel intensity values are used by humans to identify their classification, we decided on a dynamic thresholding technique to binarize the eigenvector. When examining the histogram of intensity values of the neural eigenvector (Fig. 1c), there is a large population of pixels centered around zero with a single long tail. We identified all pixels that were unique to the long tail by excluding all values that lie within range of the shorter gaussian tail. From these binarized masks, morphometrics of each primary region of the component can then be quantified, such as the axis lengths or eccentricity of the shape.

We characterized temporal dynamics of each component by extracting features from the corresponding temporal fluctuations and their frequency analysis (fig. 1d). The relative intensity fluctuations allows us to pull out temporal features of each component, like standard deviation and global maxima/minima of each component contribution. We performed wavelet analysis on these time series to characterize only highly significant frequencies (fig. S1). We calculated a power signal to noise ratio (PNR) with the 95% quantile of red noise defined by the autocorrelation value of each time series. With this ratio, significant frequencies resulted in a value above 1.

After extracting these metrics, we then compared the diverse populations of signal and artifact components, separated between vascular and other, for each feature of interest (fig. 1e−h). A full list of all metrics their respective definitions is provided, organizing between spatial, morphometric, temporal, and frequency features (fig. S2). While the data shows trends, there is not one single metric that alone could predict the classification of artifacts and signal components.

Control neural components are not distinct globular regions as those from the GCaMP line (fig. S3), but rather had co−activity with vascular units in the center of its domain. This resulted in the thresholded region to be more similar to the vascular artifacts seen in GCaMP components. However, we were still able to find example components that only had vascular spatial representation without the surrounding tissue activation. Finally, we found similar artifacts of the *other* category in the control data that are also present in the GCaMP sets of independent components.

### Characterization of spatiotemporal metrics

To investigate how well these metrics captured features of each component, we explored the coverage of the cortical surface with regions identified by the dynamic thresholding technique. Plotting all the contours of one experiment, the major footprint of the component shows the representative space of each component within brain region (fig. 2a). The majority of defined brain regions are represented by the GCaMP component footprints, with varying amounts of overlap associated with different cortical areas. Control data resulted in sparsely mapped footprints across the cortex. Further, the mapped centroid location of all components that had a thresholded region from all GCaMP experiments shows the neural components have high densities in the sensory portions of the brain (fig. 2b).

**Figure 2:**
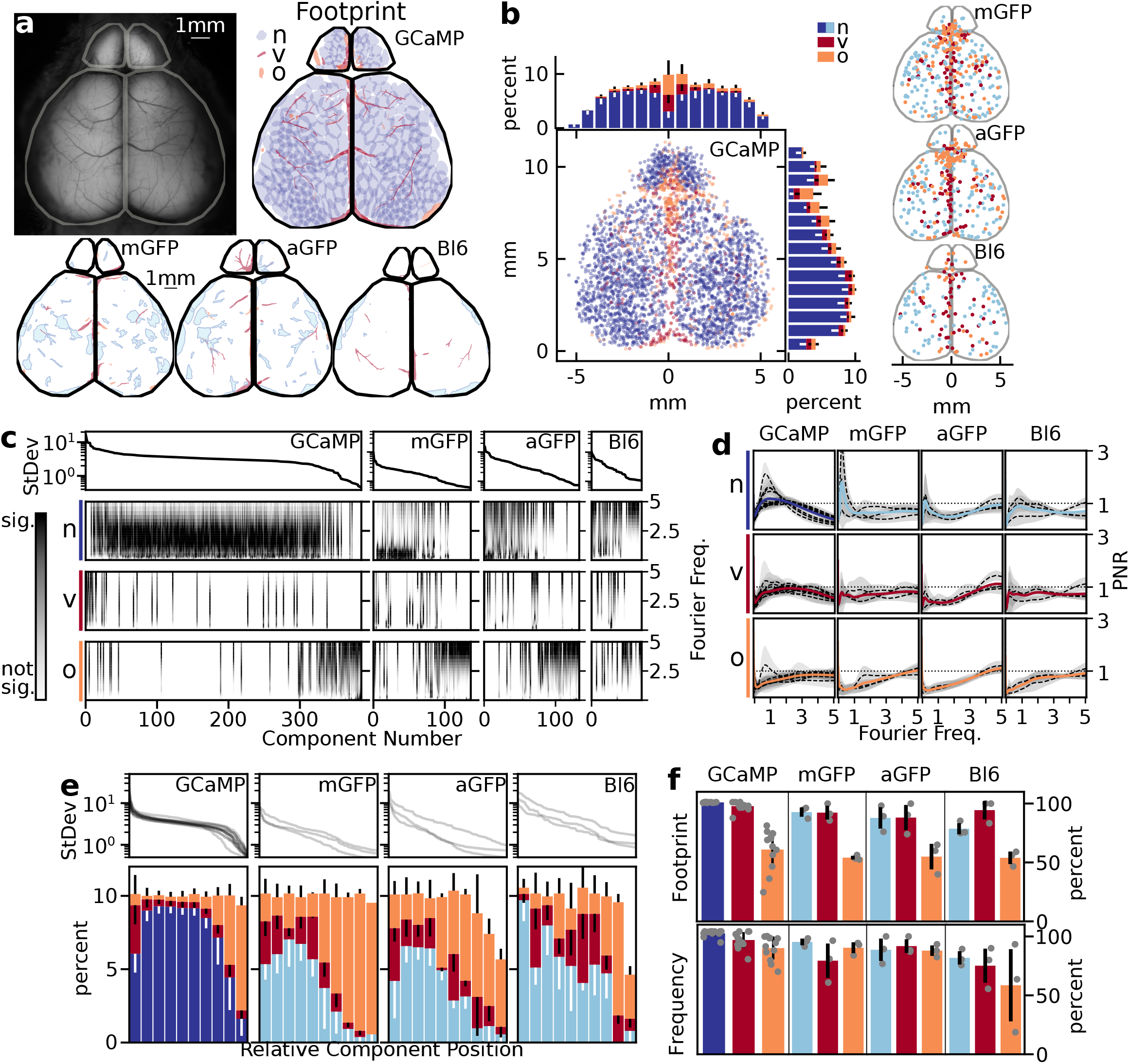
Spatial thresholding and frequency data reliably produce neural metrics. (a) Individual experiment preparation with corresponding spatial footprints by class of component: GCaMP neural (dark blue), control neural (light blue), vascular (red), other (orange). (b) All model experiments (N=12) with corresponding centroid location of each of the class of metrics. Histograms show resulting average distribution of spatial location across the field of view (error bars are standard deviation between experiments). (c) Individual experiment (same as (a)), where components are sorted by temporal variance. PNR mapped to each component and organized between the classification of component. (d) Main frequencies seen in each component class between each experimental condition. Dotted lines represent the mean within each animal, where the gray around the mean corresponds to the standard deviation of that animal. The color line corresponds to the grand mean all between experiments. (e) Relative position of the types of components between experiments and transgenic model, shown as the average and standard deviation between experiments. (f) The percent components that had footprints and frequency data that was above the noise cut−off, separated by component type and experimental condition.

Thresholded GCaMP neural components have high densities in the olfactory bulbs and posterolateral portions of the cortex, including visual, auditory, and somatosensory systems. There are less dense localization of centroids along the anteromedial portions of the cortex, including motor and retrosplenial cortices. Further, in both the GCaMP and control mice, we see the majority of artifact components localize along anatomical brain vasculature. The major venous systems including the rostral rhinal vein, the superior sagittal sinus, and the transverse sinus show high densities of artifact centroid locations ^12^. The cerebral arteries are less consistent in localizing the primary domain of their respective components. We see that many of the other artifacts align with the sagittal and lambda cranial sutures ^13^.

We investigated the effects of wavelet analysis on feature generation, by sorting each component within each experiment to its temporal standard deviation value. We then ordered each class of components based on variance and displayed a grayscale heatmap of the significant frequencies in each component across experimental conditions (fig. 2c). Taking the average of each of the global wavelet spectrum across each experiment highlighted the prominent frequencies seen in each classification (fig. 2d). Prominent GCaMP frequencies are between 0.3 to 3.5 hz, where control dynamics are seen between 0−1hz. Vascular components tend to have the same frequencies of their neural counter parts. Other components typically have faster frequencies (above 3 hz), most likely due to motion artifacts.

We looked at the overall distribution of the class of components across the relative variance position in the sorted index (fig. 2e). We found significant shifts in distribution in the types of components based on variance. Neural component were found with high variance, however they significantly tapered off nearing the noise floor. We found the highest percentage of vascular components with high variance, followed by constant low probability throughout the relative variance. The other components had the highest probability close to the noise floor, with low frequency throughout the rest of the relative variance position.

We found that wavelet analysis between all GCaMP experiments showed 95.8 *±* 3.8% of the components had significant frequencies to assess. We found that 98.4 ± 2.5% of all GCaMP neural components had significant frequencies (fig. 2f). Vascular artifacts had 93.0 *±* 6.5% and other artifacts had 86.6*±* 9.4% of components with significant wavelet frequencies within their respective class of components.

Among all GCaMP experiments, 92.3 *±* 5.6% of the components had a thresholded region to assess. However, when we looked at the breakdown of the class of components, we found that 99.8 *±* 0.2% of all neural components had a thresholded region (fig. 2e). The artifacts had fewer thresholded components, specifically the other classification; vascular artifacts had 96.6 *±* 3.6% and other artifacts had 60.2 *±* 16.2% within their respective class of components had a thresholded footprint. Of note, we found that components with low variance close to the noise threshold had increased probability of not having a domain threshold.

Overall, this indicated that all metrics can be generated for the vast majority of neural signal and vascular artifacts. However, on average, 40% of the other artifacts do not have morphometrics for their components and 13% are lacking significant frequencies. Further, this also shows changes in the relative spatial and temporal variance distributions of each of these components. The neural and vascular components align with known and predictive anatomy and were found primarily in mapped locations of increase variance. The majority other components have less variance, and therefore less contribution to the original dataset and, of the ones that had a footprint, were typically found along cranial sutures. All control data had fewer footprint and frequency metrics (fig. 2 a,c).

### Feature Selection

To build a classifier, we need to identify which metrics could distinguish between neural and artifact components. Correlation of metrics between neural signal and their respective t−statistics identified which features were most likely useful to classify a component (fig. 3a). For this process, we randomly selected seven animals from our twelve experiment dataset for determining features for training the classifier. The remaining five were used as the novel dataset for validating the machine learning performance.

**Figure 3:**
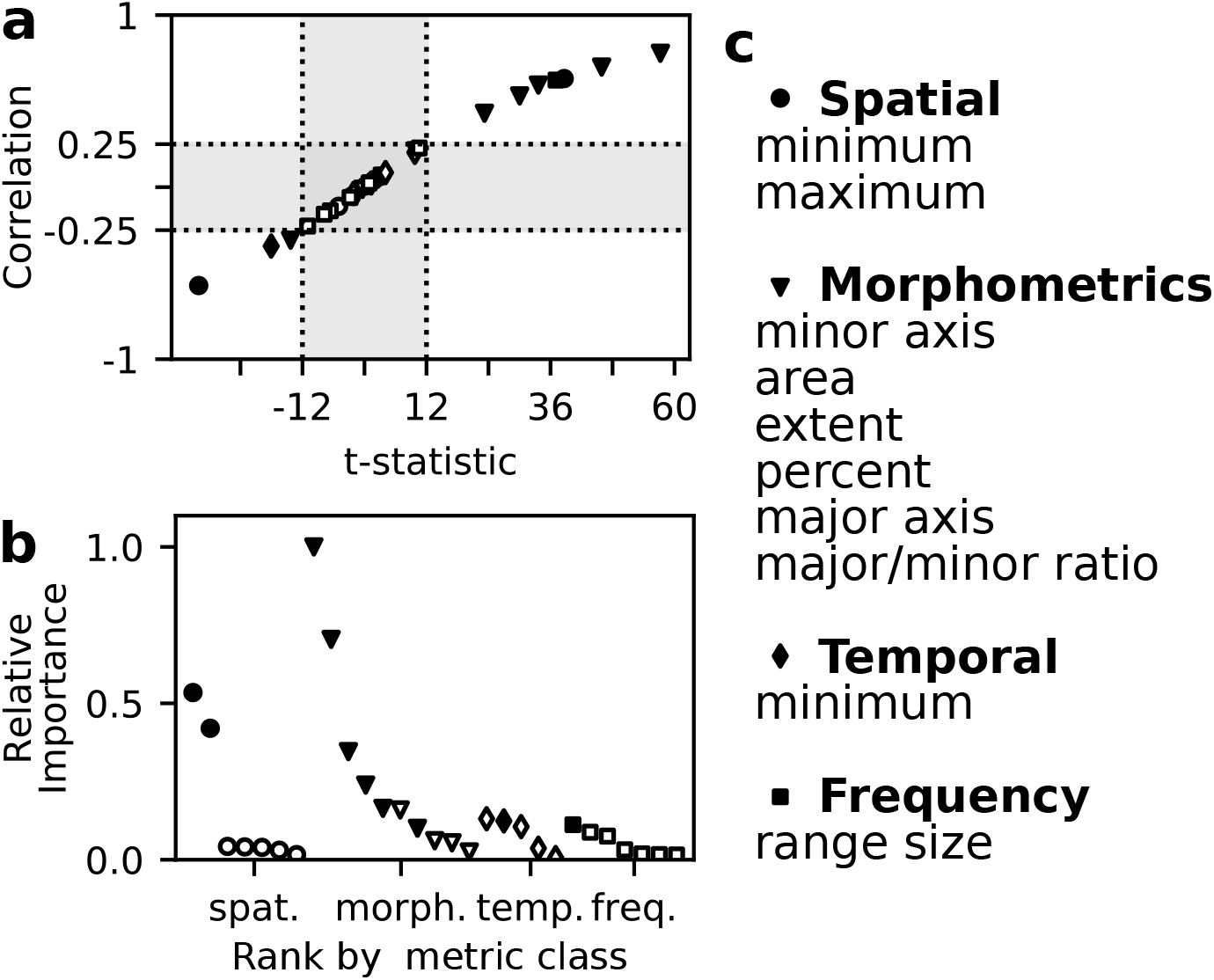
Spatial and morphological metrics are most important to classify components. (a) Correlation and t−statistic between artifact and neural components for each feature (N=7, n=2190). Spatial (circles), morphological (triangle), temporal (diamond), and frequency (square) metrics plotted. Cut off values that helped in the selection process are dotted lines, rejected values in gray. Closed points are components that met requirements. (b) Relative importance metric from the Random forest classifier plotted against each metric by their respective classes. (c) Selected metrics shown in the list within each type of feature, sorted by greatest t−statistics magnitude.

We trained the random forest classifier with all metrics to identify the importance of each feature (fig. 3b). The features having the greatest t−statistic magnitude had the most importance for proper classification. In particular, spatial and morphological metrics were found to have the highest relative importance for component classification. The final list of 10 feature metrics utilized in the machine learning process are shown (fig. 3c).

### Machine Learning Performance

We utilized the common approach of hiding a portion of the data from the learning algorithm to validate efficiency of machine learning and establish hyperparameters (fig. 4a top). Stratified sorting was used to ensure an equal ratio of artifact and neural signal was placed into each subset. We sampled and trained the random forest classifier 1000 times to see the distribution of results. We found that it performed well by all metrics assessed: mean accuracy of 97.1%, mean precision of 98.4%, and mean recall of 97.6% (fig. 4b).

**Figure 4:**
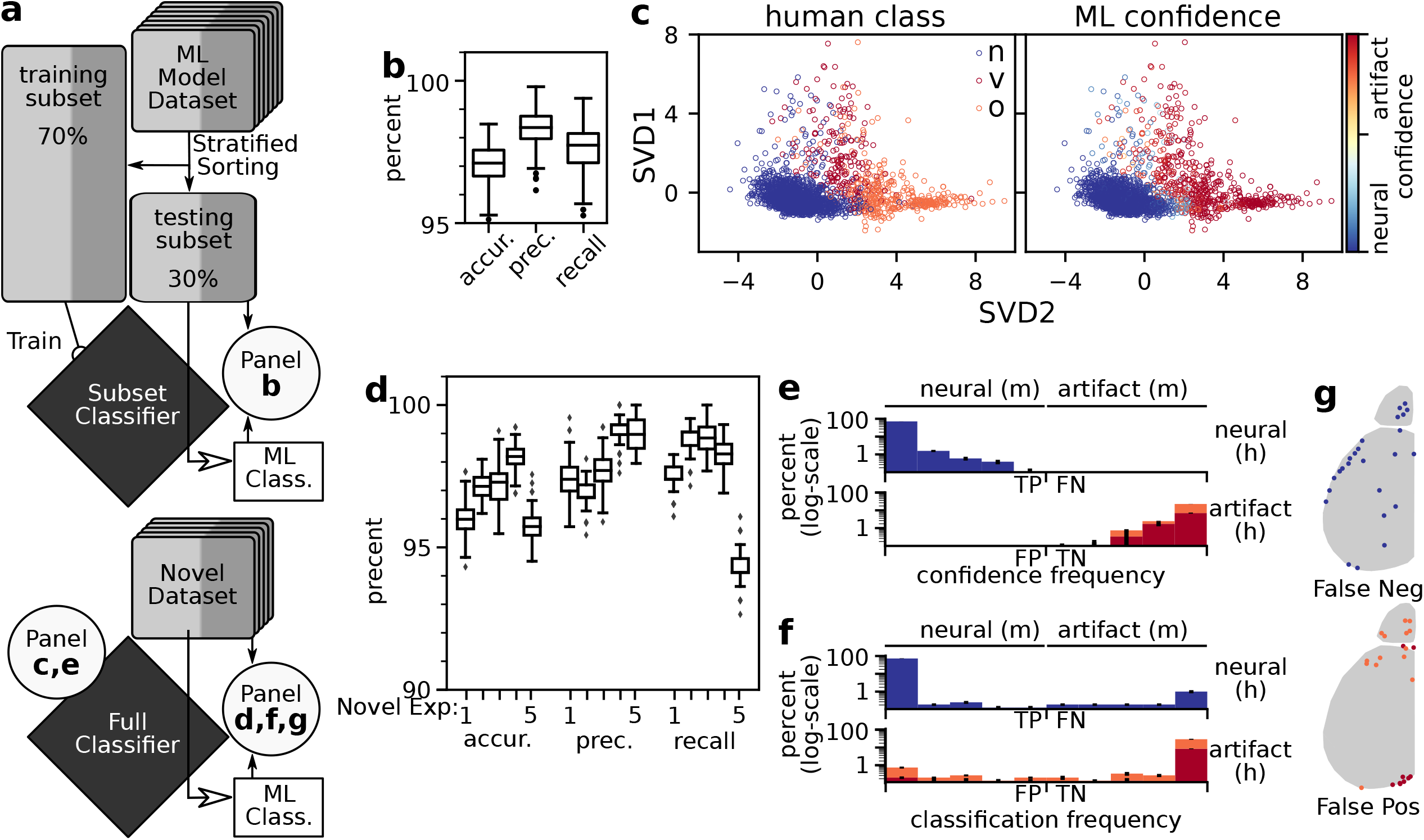
Classifier is 97% accurate on new experimental data. (a) The dataset was parsed into ML modeling dataset (N=7, n=2190) that was used to establish the machine learning pipeline and a novel dataset (N=5, n=1661) of full experiments that will not influence the classifier. Modeling data was stratified 70/30 split based on classification. 1000 iteration of training the machine learning classifier on selected metrics and validating the machine classification with human classifications. 1000 iterations of training on the full ML building dataset was performed and the novel dataset was assessed on its performance. (b) Performance of the ML training, using subsets of the ML modeling dataset. 1000 iterations resulted in accuracy, precision and recall boxplots. (c) SVD projection of metric data with human classification mapping (left) and the confidence of the ML classifier (right). (d) Performance of each of the novel datasets the full classifier’s performance, animals plotted separately showing distribution of the 1000 different trained classifiers. (e) Histogram of human classification with the percent of in each bin confidence bin (same data as c, right). Log−scale was used to highlight the low percentage points. (f) Histogram of human classification with the percent in each binned novel classification. Classification bins based on the percent each classification occurred correctly in the 1000 trials. (g) Approximate location of false negative and positives from novel datasets. True positive (TP), False positive (FP), False negative (FN), True Negative (TN), human (h), machine (m)

After establishing the efficacy of the classifier, we set out to assess the full classifier based on all data points from the machine learning dataset. We projected all features onto the first two components of a singular value decomposition (SVD), mapping both the human classification and the mean classifier confidence for 1000 iterations (fig. 4c). As expected, we saw distinct neural and artifact clusters in feature space. Interestingly, the two different types of artifacts also separated into distinct portions of the projected feature space. The confidence of the classifier showed very few components between the extremes, illustrated by the top binned confidence value distributions for each human classification (fig. 4e).

We found 71.2 *±* 0.2% of components were binned in highly confident values for neural signals (left−most bin), and 22.7 *±* 0.2 were binned in highly confident values for artifacts (right−most bin) (7.0 *±* 0.2% for vascular; 15.7 *±* 0.1% for other). This indicates that the classifier exhibits reliable confidence in the decision boundaries.

To assess the efficacy of this classifier, we then tested 1000 iterations of novel data, completely new experiments that were not involved in training the classifier (fig. 4a bottom). We plotted the resulting 1000 iterations of each experiment separately (fig. 4d). Notably, we found that the overall results were about the same as the subset classifier: mean accuracy of 96.9%, mean precision of 98.0%, and mean recall of 97.6%. From the histogram of classification frequency, we found similar results to the confidence of the classifier (fig. 4f). Among all components, 69.7 *±* 2.0 were confidently classified by machine learning as neural signal (left−most bin), where 25.8 *±* 1.4% were confidently classified as artifact (right−most bin; 7.5 *±* 0.4% of vascular; 18.2 *±* 1.6% of other). The remaining 5% either resided outside the two extreme bins or were mis−classified. We investigated the locations of the components that consistently showed false positive or false negative (fig. 4g). The majority of these components were either on the edge of the region of interest for the cortical hemispheres or within the olfactory bulb.

### Analysis of artifact signals in global mean

Removal of artifact components will ensure that neural signals are the dominant signal to be analyzed; however, during reconstruction of the data, re−addition of the global mean must occur. Thus we examined the influence that removal of artifact components has on the global mean and how filtration of the global mean should be considered. For example, vascular artifacts associated with the superior sagittal sinus contribute to the global mean and increase the range of signals recorded during periods of motion. Assessment of the global mean from GFP control experiments showed pronounced signal in these slower frequency oscillations, suggesting the use of a highpass filter fig. S4. Indeed, we found that application of a high−pass filter with a 0.5 Hz cutoff minimizes these types of global slow oscillations (fig. S5). This type of filtration should not be applied to each component individually, as there are regional networks reliant on these slower oscillations ^14^. Removal of these low frequencies from the global mean gave improved identification of the cortical patch signal sources that contribute to neural activation.

## Discussion

We have implemented an automated classification pipeline for selection of the maximally independent neural components in widefield calcium imaging videos of baseline neocortical dynamics. We identified combinations of component features that distinguish neural and artifact signal sources from decompositions of sufficient spatiotemporal sampling and feature extraction in calcium flux videos. Vascular network artifact components were found to exist independently of the globular neural signals (Figure 1) ^11^. Vascular networks exhibited dynamics within similar frequency ranges as neural components, due to the refraction or occlusion of the GCaMP signal by the vasculature. Given that vascular artifacts had overlapping temporal and frequency metrics with the neural components, we found that a set of spatial and morphological feature metrics gave optimal classifier performance during machine assignment of components to different signal classes. Furthermore, we showed another class of movement and optical artifacts that are present in both control and experimental GCaMP recordings. Frequency analysis from these other artifacts had higher power in the faster dynamics (above 3 Hz) and are due to motion. Indeed many of these other artifacts were associated with vascular tissue displacments during animal movements. Thus, it is therefore necessary to removed these other artifacts during video pre−processing. Similar approaches have been implemented in fMRI studies, where researchers built an ICA−based classifier to scrub motion artifacts ^15, 16^.

The data rebuild of identified neural components with mean filtration is a statistically valid process for isolating neural signals. Our recordings consisted of large sets of densely sampled image frames having at least 12 bits of dynamic range across pixel intensities. We hypothesize that these sampling conditions coupled with a strong neuronal GCaMP signal−to−noise ratio optimizes ICA’s signal de−mixing ability to isolate functionally discrete patches of cerebral cortex from other physiological signals. In control recordings lacking a calcium sensor to report neuronal dynamics, high quality isolation of signal components was not attained given equivalent video sampling conditions. The dynamics of vascular and neural tissues are energetically and thus physiologically linked and the interplay between the hemodynamic responses and neural signals is known ^10, 17^. Even so, we found that neural GCaMP components comprise discrete units across the cortex (Figure 2) ^11^. In tissue expressing a contrast agent, such as GFP, the optical hemodynamics are enhanced and result in widespread regional effects among the cerebral lobes from control animals. Wavelet analysis on the global mean and individual neural components show the dominant signals extracted from GCaMP animals as being in a faster frequency range than cortical hemodynamics (*>*1 Hz). Our results indicate that neural GCaMP signals heavily outweigh the neocortical hemodynamic signals in decomposed independent components of densely sampled wide−field calcium imaging videos.

Functional imaging in unanesthetized, behaving animals gives insight into the nature of physiological processes, however, nontrivial challenges arise during such sessions with intermixed sets of time varying signals. This work demonstrates that signal components having maximal statistical independence captured in sufficiently sampled mono−chromatic calcium flux videos exhibit a combination of spatiotemporal features that allow machine classification of signal type. Implementation of automated machine classifiers for neural signals is practical given densely captured arrays of spatially and temporally variant data gathered from individual subjects.

## Figures

**Figure S1:**
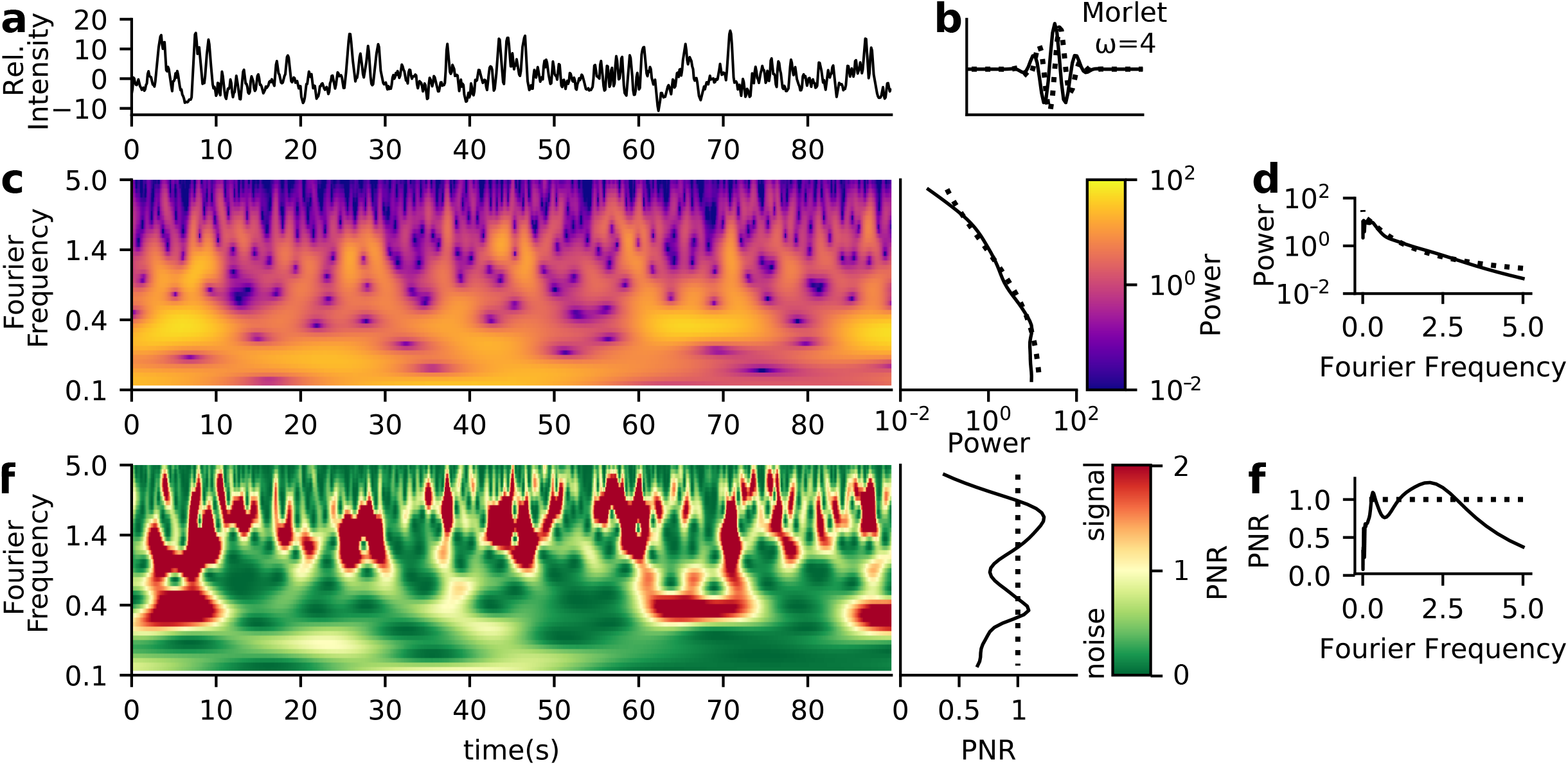
Morlet wavelet transform can be used to create a signal to noise ratio that indicates frequencies of high likelihood of signal. (a) Example neural time series, 90 sec of data recorded at 10hz reported in the temporal portion of a component (b) Morlet wavelet (=4) was used for the wavelet transform. (c) The power spectra of the wavelet transform (colorbar, purple to yellow) and the global spectral analysis (black, right). The 95% quantile is shown in dashed lines on the global spectral analysis. Reformatting the frequency spacing, produces (d). (d) A signal to noise power ratio is calculated by dividing the power spectra by the 95% quantile. All values above 1 (dashed line) would indicate a high probability of signal. Anything below 1 would most likely be considered noise (colorbar, green to red). Reformatting the frequency spacing, produces (f).

**Figure S2:**
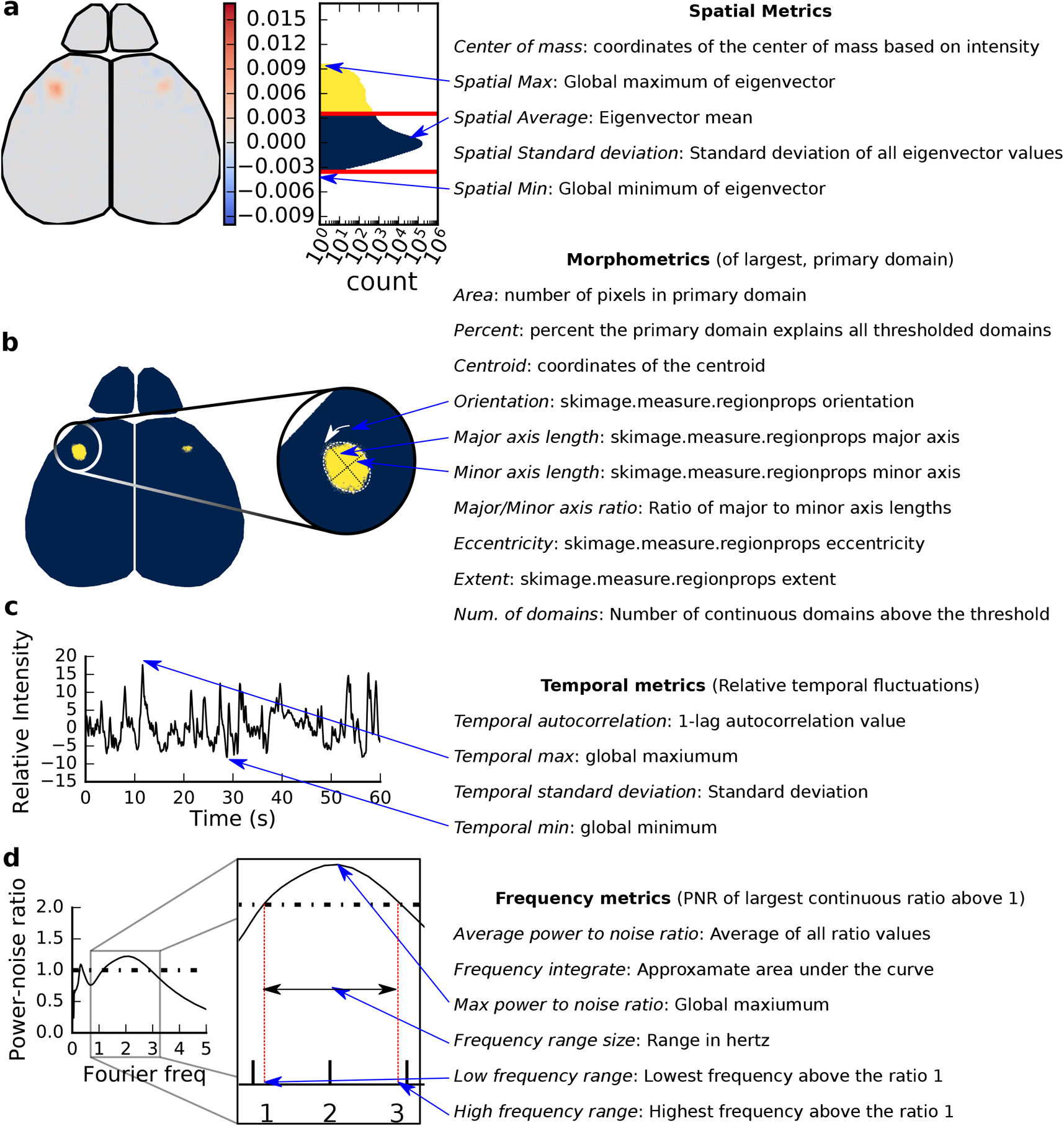
Spatial, Morphometric, Temporal, and Frequency features extracted from components. (a) Spatial metrics from statistical characteristics of each eigenvector (spatial representation of the component). The histogram of all eigenvector values is shown the right of the eigenvector. (b) Morphometrics collected from the binarized thresholded masked region of the eigenvector. The largest (primary) domain was used to generate the features for each eigenvector. The majority of metrics calculated utilizes sci−kit image region properties. (c) Temporal metrics are statistical descriptors from the corresponding row of the mixing matrix for each eigenvector. (d) Frequency analysis was done on the mixing matrix row, utilizing the PNR calculated from wavelet analysis (Figure S1). The longest of all continuous frequencies was used to extract each feature.

**Figure S3:**
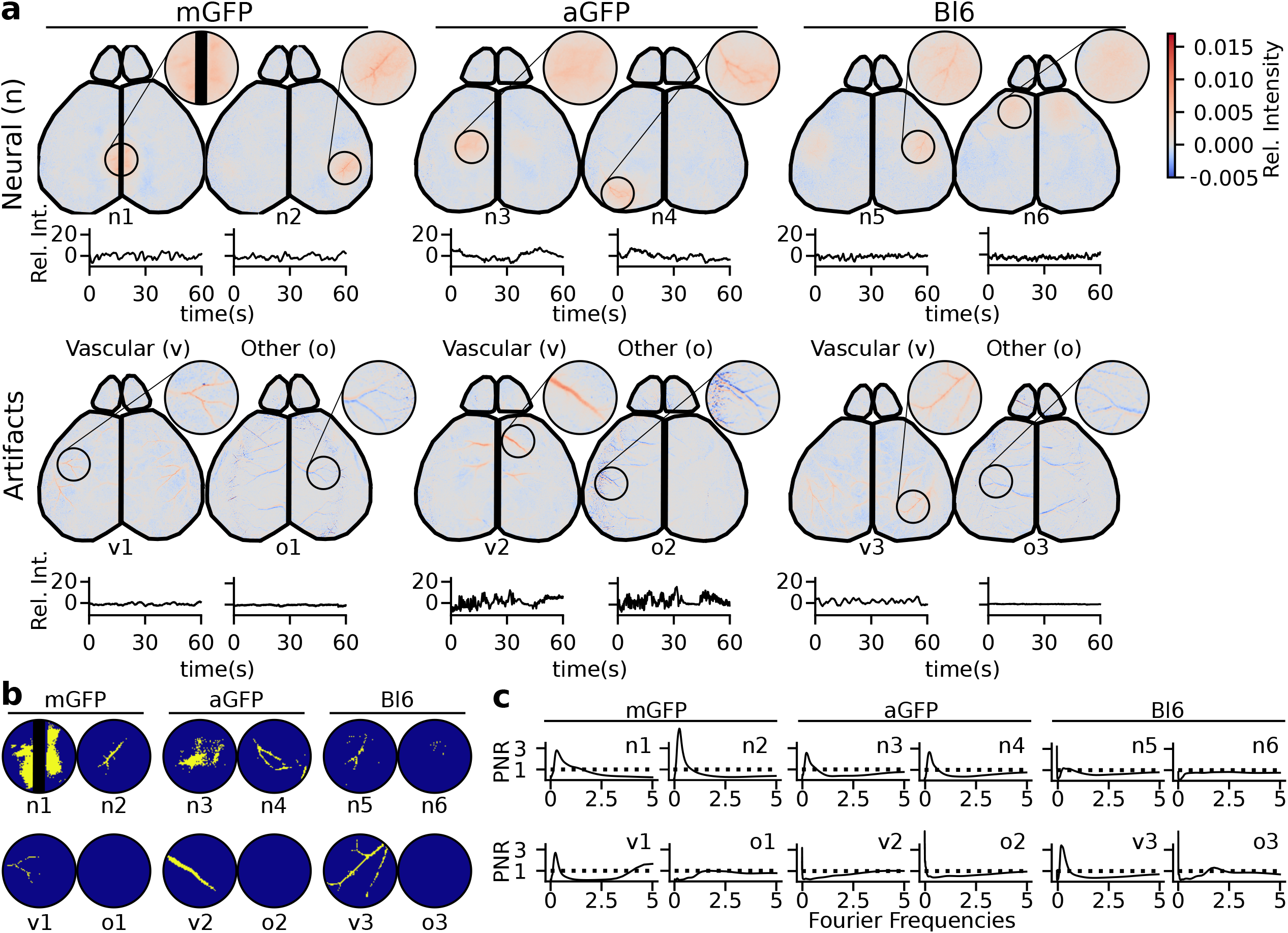
Examples of control components, resulting in similar artifact components to GCaMP recordings. Control components from 20 minutes of recording from cx3cr1 GFP (microglia; mGFP, left), adlh1 GFP (astrocyte; aGFP, center), and Black 6 (Non−transgenic; Bl6, right) mice. Two IC examples from each control group corresponding to hemodynamics/neural activity (top) and artifacts (bottom). Artifacts chosen shows a vascular and other artifact commonly seen in GCaMP recordings. Similar data description in regards to temporal and spatial representations as seen in Figure 1. (b) Examples of control binarization of the eigenvector only showing the windowed spatial representation on the key portions of the eigenvector. (d) Examples of neural and artifact wavelet analysis shown in the to signal power−noise ratio (PNR) plots.

**Figure S4:**
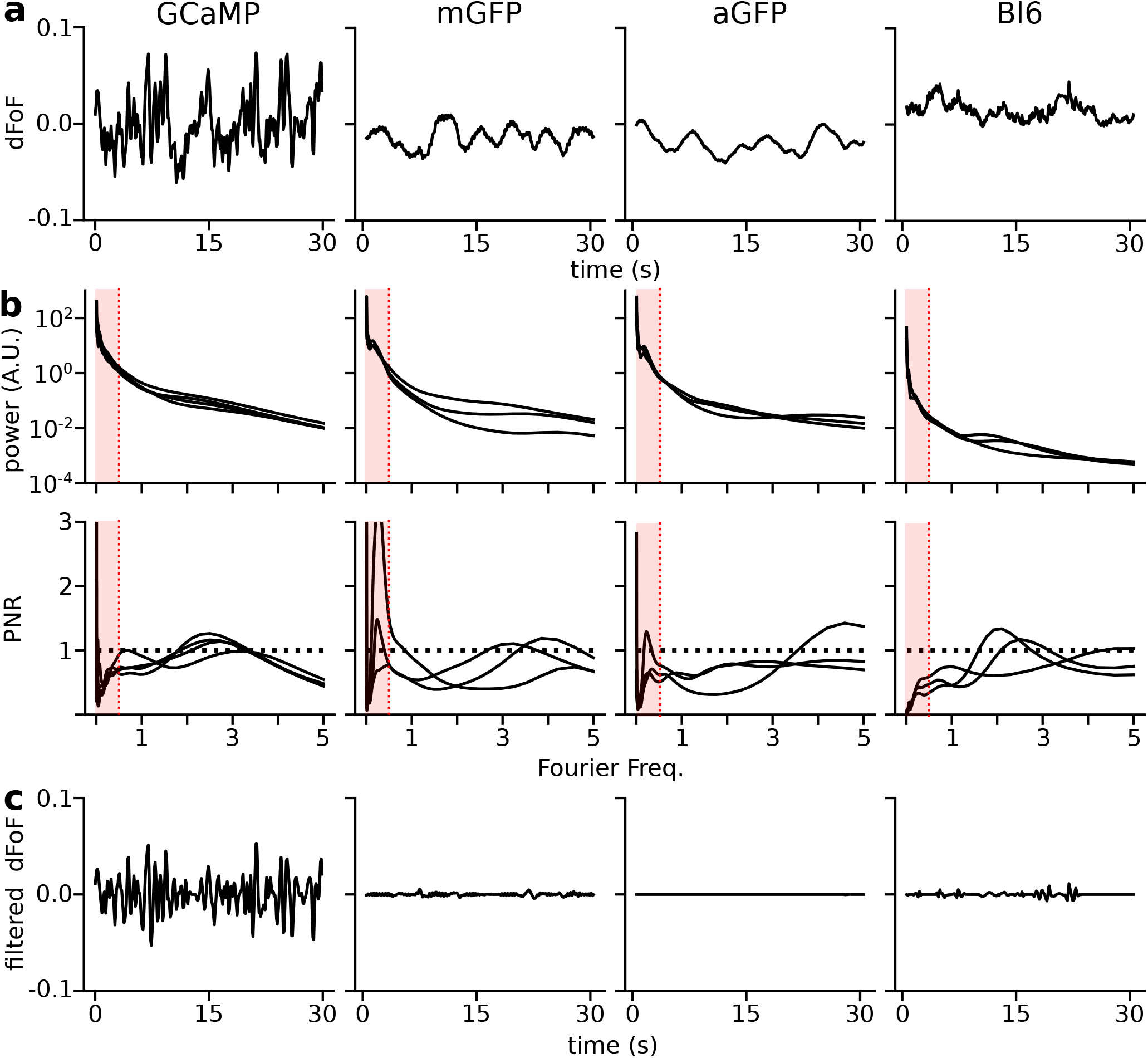
Mean filtration to minimize global slow oscillations seen in GFP control data. (a) 30sec examples of the global mean that was subtracted and stored at the initiation of the pipeline, before the decomposition into eigenvectors for GCaMP, mGFP, aGFP and Bl6. (b) Global wavelet spectrum (top) and its corresponding power to noise ratio (PNR; bottom) of GCaMP (N=4), mGFP (N=3), aGFP (N=3), and Bl6 (N=3), red indicates the omitted frequencies from our applied high− pass filter. (c) High−pass filtration results of the same 30 sec in (a).

**Figure S5:**
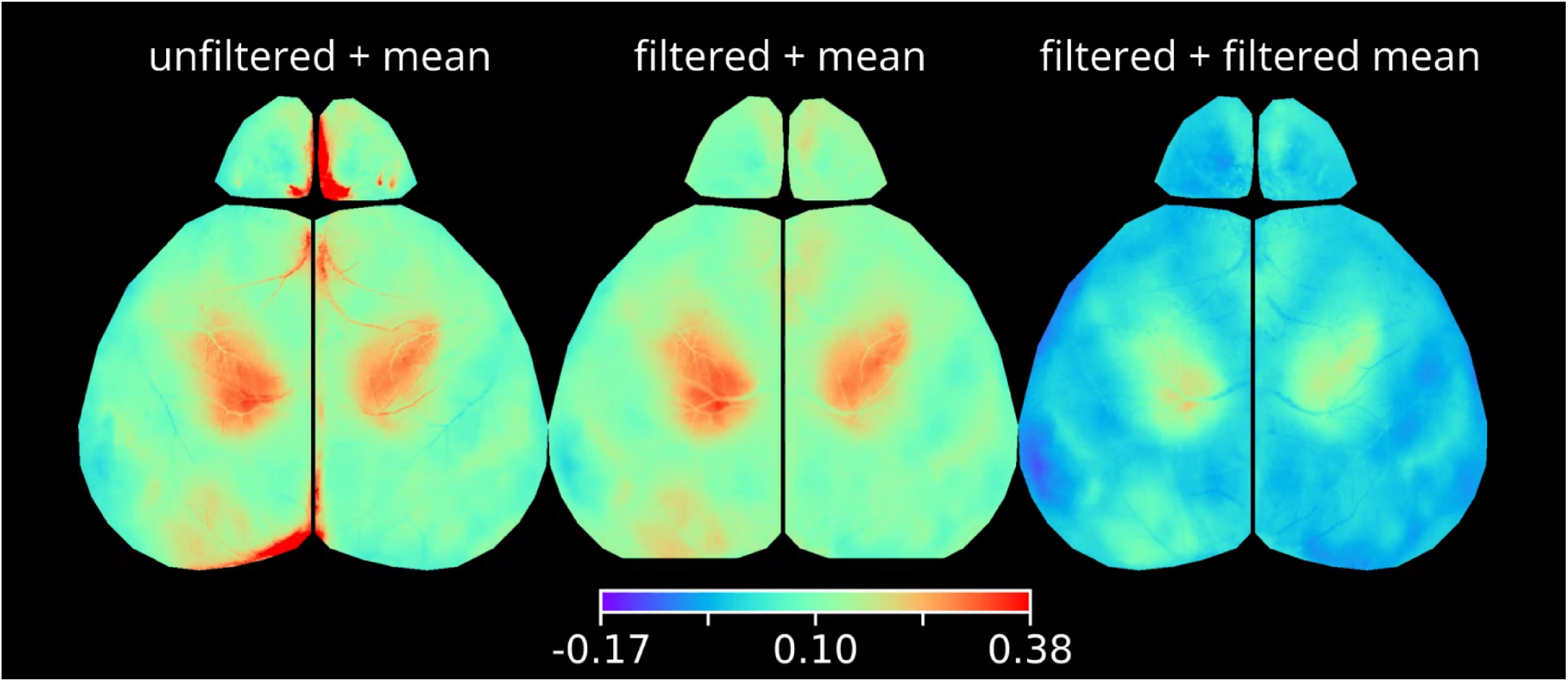
Readdition of the global mean with high−pass filter better represents neural signal. Rebuilt video from all components (left) or with only neural components (center, right) with mean re−addition from the original global mean (left, center) or the global mean with 0.5 Hz high pass filtration (right). All movies are on the same scale of change in fluorescence over mean fluorescence (rainbow colorbar)

## Methods

### Mice

All animal studies were conducted in accordance with the UCSC Office of Animal Research Oversight and Institutional Animal Care and Use Committee protocols. P21−22 Snap25 GCaMP6s transgenic mice (JAX: 025111), Cx3cr1 GFP (JAX: 005582), and Aldh1 GFP (MGI: 3843271) were maintained on a C57/Bl6 background in UCSCs mouse facilities. To identify Snap25 GCaMP expressing mice, a single common forward primer (5’−CCC AGT TGA GAT TGG AAA GTG−3’) was used in conjunction with either transgene specific reverse primer (5’−ACT TCG CAC AGG ATC CAA GA−3’; 230 band size) or control reverse primer (5’−CTG GTT TTG TTG GAA TCA GC−3’; 498 band size). The expression of this transgene resulted in pan−neuronal expression of GCaMP6s throughout the nervous system. To identify GFP expressing mice a forward (5’−CCT ACG GCG TGC AGT GCT TCA GC−3’) and reverse (5’−CGG CGA GCT GCA CGC TGC GTC CTC−3’; 400 band size) PCR amplification was used to identify which animals had the GFP transgene. At the end of each recording session, the animal was either euthanized or perfused and the brain dissected.

7 animals used in this study were experiment and control mice to study perinatal penicillin exposure effects on cerebral networks.^18^ These methods work independent of experimental condition and the perinatal penicillin had little effect on domain parcellation.

### Surgical procedure

All mice were anesthetized with isoflurane (2.5% in pure oxygen) for the procedure. Body temperature was maintained at 30C for the duration of the surgery and recovery using a feedback−regulated heading pad. Lidocaine (1%) was applied subcutaneous on the scalp, followed by careful removal of skin above the skull. Opthalmic ointment was used protect the eyes during the surgery. The cranium was attached using to two head bars using cyanoacrylate, one across the occipital bone of the skull and the other on the lateral parietal bone.

After the surgery was complete, mice were transferred to a rotating disk for the duration of the recording. At the end of the recording session, the animal was euthanized and the brain dissected.

### Recording calcium dynamics

In−vivo wide−field fluorescence recordings were collected in a minimally invasive manner. Imaging through the skull by single−photon excitation light from two blue LED light (470 nm; Thorlabs M470L3) produces a green fluorescent signal that is collected through coupled 50*mm* Nikon lenses (f5.6 / f1.2, optical magnification *∼* 1x) into a scientific CMOS camera (PCO Edge 5.5MP, 6.5*μ*m pixel resolution). The top lens in the tandem lens setup was used to focus on the cortical surface, thereby lowering the magnification slightly; anatomical representation for each pixel corresponded to 6.9 *±* 0.2*μ*m (min: 6.7*μ*m, max: 7.2*μ*m). Excitation light was filtered with a 480/30 nm bandpass (Chroma Technology AT480/30x) and the emission signal was filtered with 520/36 nm bandpass (Edmund Optics 67−044). Data collection was performed in a dark, quiet room with minimal changes in ambient light or sound, thus the brain activity recorded was resting state without direct stimulation. Raw data was written directly as a set of 16 bit multi−image TIFF files.

The total amount of data recorded for each animal was generally at least 40 min and the amount of time in between video segments was less than 1 minute. All analyzed data consisted of two sequential full spatial resolution recordings concatenated together giving 20 min of data sampled at 10 frames per second for each video decomposition^11^.

### Dynamic Thresholding

The spatial histogram of eigenvector values across each component can be visualized as a single tailed gaussian distribution centered around 0, where the tail represents the spatial footprint of each component. The two edges of the distribution are first identified. The boundary closer to 0 is taken as the edge of the central noise distribution, and that boundary is used to define the dynamic threshold on the opposite side of 0. Any values outside of this noise distribution and is part of the wider tail are included in the binarized domain of the component. If the tail was negative, the component was flipped spatially and temporally for visualization purposes.

### ICA and Data processing

ICA decompositions were run on a single computing cluster node having 1024 GB of RAM. After ICA processing, metric extraction was computed on local computers with 16−32GB of RAM.

### Metric generation and classification of Neural Independent Components

An ensemble random forest classifier from the scikit−learn 23 package^19^ was used to train and classify between human scored signal and artifact components ^20^, based on features calculated from each component.

### Wavelet Mean filtration

Wavelet decomposition on the time series signals were performed with a ω = 4 morlet wavelet family, code adapted from C. Torrence and G. Compo,^21^ available at URL: http://paos.colorado.edu/research/wavelets/ Significance was determined using the 95th percentile of a red−noise model fit to the time series autocorrelation. Frequency distributions are all displayed as the ratio of the global wavelet specturm, relative to the noise cutoff. For wavelet filtering, the original signal was rebuilt excluding all frequency signals in a certain range.

### Statistical significance

Statistical significance was calculated using OLS models from `statsmodel.formula.api` with Holm−Sidak multiple testing correction (*p* 0.5: *; *p* 0.01: **; *p* 0.001: ***). Model significance is determined by the F−statistic, and significance of two−group analyses (p*>*| t |) are calculated with t−tests.

## Acknowledgements

The authors acknowledge C. Santo Thomas for maintaining the lab mouse lines, and University of California Santa Cruzs Hummingbird Computational Cluster for support and node maintenance. This work was supported by Startup funds from University of California, Santa Cruz, Division of Physical and Biological Sciences, grants from the National Institutes of Health, USA (NIH T32 GM 133391) to S.C.W. and B.R.M, and by a Hellman Fellows Fund Award to J.B.A. Funding for D.A. was provided by the UCSC Maximizing Access to Research Careers (MARC) program (T32−GM007910) and the UCSC Initiative for Maximizing Student Development (IMSD) (R25−GM058903).

## Contributions

ICA filtering code was written by S.C.W. All recordings, metric extractions, mean frequency analysis, feature extraction and analysis, machine learning pipeline were performed by B.R.M. Optimizing and determination of hyperparameters was done by D.A. J.B.A. oversaw the project and provided feedback to experimental design, results, and paper preparation. The manuscript was prepared by B.R.M., with input from all authors.

